# Dynamic interplay between reward and voluntary attention determines stimulus processing in visual cortex

**DOI:** 10.1101/2021.06.17.448788

**Authors:** Ivan Grahek, Antonio Schettino, Ernst H.W. Koster, Søren K. Andersen

**Affiliations:** Department of Cognitive, Linguistic, & Psychological Sciences and Carney Institute for Brain Science, Brown University, Providence, RI 02912, USA; Department of Experimental Clinical and Health Psychology, Ghent University, Henri Dunantlaan 2, B-9000, Ghent, Belgium; Erasmus Research Services, Erasmus University Rotterdam, Burgemeester Oudlaan 50, 3062 PA, Rotterdam, The Netherlands; Institute for Globally Distributed Open Research and Education (IGDORE); School of Psychology, University of Aberdeen, William Guild Building, Aberdeen, AB24 3FX, United Kingdom

**Keywords:** voluntary attention, attentional control, reward, motivation, EEG, feature-based attention, steady-state visual evoked potentials, frequency tagging, Bayesian multilevel modeling

## Abstract

Reward enhances stimulus processing in the visual cortex, but the mechanisms through which this effect occurs remain unclear. Reward prospect can both increase the deployment of voluntary attention and increase the salience of previously neutral stimuli. In this study we orthogonally manipulated reward and voluntary attention while human participants performed a global motion detection task. We recorded steady-state visual evoked potentials (SSVEPs) to simultaneously measure the processing of attended and unattended stimuli linked to different reward probabilities, as they compete for attentional resources. The processing of the high rewarded feature was enhanced independently of voluntary attention, but this gain diminished once rewards were no longer available. Neither the voluntary attention nor the salience account alone can fully explain these results. Instead, we propose how these two accounts can be integrated to allow for the flexible balance between reward-driven increase in salience and voluntary attention.

## Introduction

Maximizing rewards and avoiding punishments are among the main determinants of human behavior. In order to increase the probability of obtaining a reward, reward-related information needs to be prioritized. Selective attention is crucial for adaptive behavior as it facilitates the processing of relevant over irrelevant information in the environment (Chun, Golomb, & Turk-Browne, 2011; Desimone & Duncan, 1995). This process depends on our current goals (e.g., looking for car keys in the living room) and salience of stimuli (e.g., a loud noise; Corbetta & Shulman, 2002; Posner, 1980; Theeuwes, 2010). Recent research has indicated that motivation can influence selective attention by impacting both of these factors. Reward expectation can enhance voluntary selective attention, and reward associations can change the salience of previously neutral stimuli. In most situations, attention is guided by the combination of both voluntary allocation of attention and reward history of stimuli (Awh, Belopolsky, & Theeuwes, 2012). For example, while we are searching for keys (goal-relevant target) our attention can be captured by a cake (goal-irrelevant distractor). These two ways in which rewards influence selective attention have been commonly studied in isolation and the neural mechanisms through which they jointly guide attention remain unclear. Specifically, it remains unclear how voluntary selective attention and reward history interact to determine the processing of goal-relevant and irrelevant stimuli in the visual cortex.

Voluntary selective attention is enhanced when individuals anticipate that they can earn rewards for good task performance (Botvinick & Braver, 2015; Krebs & Woldorff, 2017; Pessoa, 2015). A number of fMRI and EEG studies found reward-based increases in attention in preparation for upcoming target stimuli. These studies have shown that such increases are driven by enhanced activity in frontoparietal regions involved in attentional control (Krebs, Boehler, Roberts, Song, & Woldorff, 2012; Pessoa & Engelmann, 2010; Schevernels, Krebs, Santens, Woldorff, & Boehler, 2014) and by enhanced task-set representations in these regions (Etzel, Cole, Zacks, Kay, & Braver, 2016; Wisniewski, Reverberi, Momennejad, Kahnt, & Haynes, 2015). While these studies suggest that reward influences attentional control via neuronal modulations in the frontoparietal network, it remains unclear how such modulations translate to affect the processing of attended and unattended stimuli in visual cortex.

Within a largely independent research line, a set of studies has focused on the processing of stimuli associated with earning rewards. These studies have demonstrated that stimuli currently or previously associated with rewards capture attention in an automatic fashion, even when this conflicts with current goals (Anderson, 2016; Awh, Belopolsky, & Theeuwes, 2012; Chelazzi, Perlato, Santandrea, & Della Libera, 2013; Failing & Theeuwes, 2017). Behavioral studies have demonstrated that stimuli predictive of rewards capture attention, and that they can do so in subsequent trials when rewards are no longer present (Anderson, Laurent, & Yantis, 2011; Della Libera & Chelazzi, 2009; Failing & Theeuwes, 2014). Event-related potential (ERP) studies have shown that stimuli related to rewards receive increased sensory processing, and attentional capture by rewarding stimuli can be related to changes in the early processing of such stimuli in the visual cortex (i.e., increase in the P1 ERP component; Donohue et al., 2016; Hickey, Chelazzi, & Theeuwes, 2010; Luque et al., 2017; MacLean & Giesbrecht, 2015). However, other studies have not found evidence for such early modulations in the visual cortex, and instead reported changes at later stages of stimulus processing (increased N2pc ERP component and improved decoding in later processing stages; Qi et al., 2013; Tankelevitch et al., 2020). Similarly, fMRI studies have also shown reward-related increases in sensory processing (Serences, 2008). More specifically, one study (Hickey & Peelen, 2015) provided evidence for the simultaneous enhancement in representation of reward-related stimuli and suppression of stimuli devoid of a specific motivational value. Using multivoxel pattern analysis and decoding technique, these authors found a gain increase in object-selective visual cortex for stimuli paired with rewards, while those not associated with this incentive were suppressed.

The reviewed findings thus point toward two mechanisms through which rewards influence selective attention. First, the prospect of earning rewards increases the voluntary allocation of attention. Second, rewards can increase the salience of previously neutral stimuli leading them to capture attention in a more automatic fashion. Importantly, the effects of reward history and voluntary attention are often difficult to disentangle, and they are often confounded in cognitive tasks (Maunsell, 2004). For example, common paradigms for studying both reward processing and attention include the association between allocating attention in a specific way (e.g. toward a location and a feature) and receiving a reward (e.g. a monetary reward, or the intrinsic reward of following the task instructions and solving the trial correctly). Further, both increases in voluntary attention and stimulus salience can lead to increased sensory gain in the visual cortex. Thus, it remains unclear which reward-related changes in stimulus processing in visual cortex occur as a consequence of voluntary selective attention, and which changes result from alterations in stimulus salience. Most importantly, reward-driven dynamic interactions between voluntary attention and changes in stimulus salience remain underexplored.

Theoretical models that focus on the relationship between incentives and attention commonly focus on either the voluntary attention or the salience aspect of their interaction. Although not mutually exclusive, these models make different predictions about the way in which rewards influence attention. One option is that rewards influence stimulus processing by increasing the amount of voluntary attention deployed toward these stimuli. This hypothesis can be derived from models that focus on the role of motivation in the allocation of attention and cognitive control (Brown & Alexander, 2017; Holroyd & McClure, 2015; Shenhav, Botvinick, & Cohen, 2013; Verguts, Vassena, & Silvetti, 2015). These models propose that the amount of attention allocated toward stimuli is dependent on the amount of rewards which are expected for doing so. Another possibility is that rewards increase stimulus salience and thus capture attention automatically, independently of voluntary attention. This view can be derived from theoretical models highlighting the role of reward history in guiding selective attention (Anderson, 2016; Awh et al., 2012; Chelazzi et al., 2013; Failing & Theeuwes, 2017). These models propose that the processing of stimuli linked to high rewards is facilitated while the processing of other stimuli is suppressed, and that this effect is long lasting, even when rewards are no longer available. Importantly, although not explicitly incorporated into the current theoretical frameworks, motivation influences both voluntary attention and changes stimulus salience. Here we sought to assess the effects of both of these mechanisms on stimulus processing in visual cortex, and in that way investigate how these two mechanisms interact to guide stimulus processing and optimize behavior.

In this study, we orthogonally manipulated voluntary attention and reward probability in order to assess how they interact within a single paradigm. To this end, we adopted an established feature-based attention paradigm (e.g., Andersen, Müller, & Hillyard, 2009; Andersen & Müller, 2010). On each trial, two superimposed random dot kinematograms (RDKs) of different color (red and blue) were presented concurrently and participants were instructed, on a trial-by-trial basis, to attend to one of them in order to detect infrequent coherent motion targets. Thus, these two RDKs served as goal-relevant (attended) and goal-irrelevant (unattended) stimuli, respectively^1^. Critically, after a baseline period used as control condition, these two colors were associated (via explicit instruction upon completion of the baseline phase) with a low or high probability of earning a reward in a training phase. We subsequently examined the influence of the previous reward history in the test phase, in which rewards were no longer available. The two RDKs flickered at different frequencies, thereby driving separate steady-state visual evoked potentials (SSVEPs). SSVEPs offer the unique advantage of simultaneously tracking the processing of multiple stimuli as the specific oscillatory response of each stimulus can be extracted (frequency tagging), and the two resulting signals can be compared to each other (Andersen & Müller, 2010; Kashiwase, Matsumiya, Kuriki, & Shioiri, 2012; Müller, Teder-Sälejärvi, & Hillyard, 1998). Voluntary attention is known to increase SSVEP amplitudes of attended stimuli (Morgan, Hansen, & Hillyard, 1996). Further, SSVEP amplitudes are highly sensitive to changes in the physical salience of stimuli and are increased for more salient stimuli (Andersen, Müller, & Martinovic, 2012). Thus, the SSVEP amplitudes capture the changes in sensory gain resulting from either the top-down influences of voluntary attention, or the bottom up changes in salience. Hence, analyzing SSVEPs in this design provided us with the ability to simultaneously track the visual processing of attended and unattended stimuli related to high or low rewards respectively. This design thus enabled us to experimentally dissociate between the effects of voluntary attention (instructions about which color to attend to) and reward probability (stimulus-reward pairings).

We tested predictions arising from the theoretical models developed to account for the effects of rewards on cognitive control (Brown & Alexander, 2017; Holroyd & McClure, 2015; Shenhav et al., 2013; Verguts et al., 2015) and the effects of reward history on attention (Anderson, 2016; Awh et al., 2012; Chelazzi et al., 2013; Failing & Theeuwes, 2017), respectively. The first class of models predict that reward influences sensory processing through voluntary attention, and the second class of models predict that rewards directly modulate stimulus salience. Both groups of models predict better behavioral performance and enhanced processing (higher SSVEP amplitudes) of the stimuli related to high rewards. However, the strict reward history view would predict that the processing of the high reward stimuli will be enhanced irrespective of voluntary attention (i.e., equally when they are unattended or attended), while the strict voluntary attention view would predict that the processing of the high reward stimuli will be enhanced only when they are attended. Finally, the reward history view predicts that these effects will persist when rewards are no longer available (in our paradigm, during the test phase), while the voluntary attention view predicts that the processing of both high and low reward stimuli will return to baseline levels. Here we tested these predictions by independently manipulating voluntary attention and reward, which allowed us to assess the contribution of each of these factors and possible interactions. Most importantly, this design allowed us to investigate how reward-driven changes in voluntary attention and reward-driven stimulus salience jointly determine stimulus processing in visual cortex leading to behavioral adaptations and increasing the amount of earned rewards.

## Methods

### Participants

We tested 48 participants with normal or corrected-to-normal vision and no history of psychiatric or neurological disorders. Four participants were excluded due to technical problems during EEG recording and one person was excluded due to noisy EEG data. Thus, the final data set consisted of 43 participants (29 females, 14 males; median age = 22). Participants received a fixed payoff of 20 €, plus up to 6 € depending on task performance (on average 25.5 €). The study was approved by the ethics committee of Ghent University.

### Stimuli and task

We used a coherent motion detection task (Andersen & Müller, 2010; *Figure 1A*), in which participants were presented with two overlapping circular RDKs of isoluminant colors (red and blue) on a grey background. Viewing distance was fixed with a chinrest at 55 cm from the 21-inch CRT screen (resolution of 1024 x 768 pixels, 120 Hz refresh rate). At the beginning of each trial, participants were instructed which of the two RDKs to attend by a verbal audio cue: “red” (241 ms) or “blue” (266 ms). The two RDKs had a diameter corresponding to 20.61 degrees of visual angle and consisted of 125 randomly and independently moving dots each (0.52 degrees of visual angle per dot). The two RDKs flickered at different frequencies: 10 Hz (6 frames on / 6 frames off) and 12 Hz (5 frames on / 5 frames off). 40% of trials contained no coherent motion intervals. The other 60% of trials contained one, two, or three coherent motion intervals, occurring with equal probability in the attended and unattended color RDK. This was done to ensure that participants maintained attention throughout the trial. During these intervals, dots in one of the RDKs moved with 75% coherence in one of four cardinal directions (up, down, left, or right) for 300 ms. The earliest onset of coherent motions was 750ms after onset of the RDKs and subsequent coherent motions within the same trial were separated by at least 600ms to allow for an unambiguous assignment of detection responses to preceding coherent motions. Participants had to detect the occurrence of coherent motion in the attended RDK as fast as possible by pressing the space key on a standard AZERTY USB keyboard while ignoring such coherent motion in the unattended RDK. Responses occurring between 275 ms and 875 ms after coherent motion onset of the attended or unattended dots were counted as hits or false alarms, respectively. Correct responses were followed by a tone (200 ms sine wave of either 800 or 1,200 Hz, counterbalanced across participants). Late or incorrect responses were followed by an error sound (200 ms square wave tone of 400 Hz).

**Figure 1.**
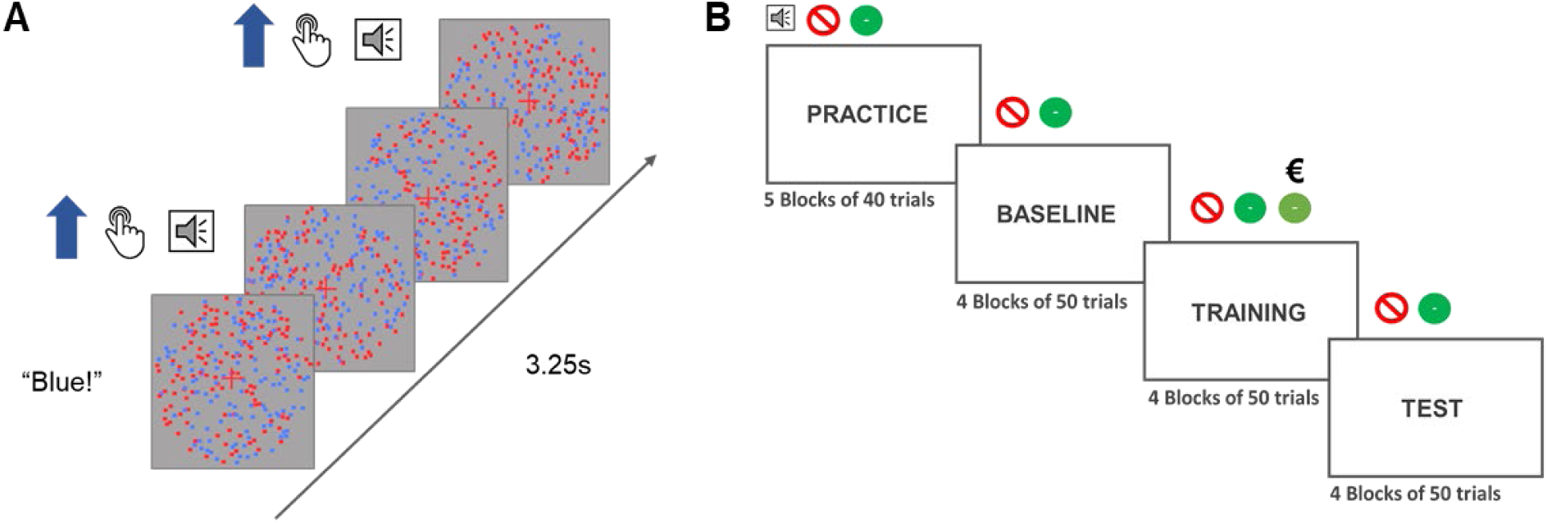
Depiction of a single trial and the phases of the experiment. **A.** Each trial started with an audio cue (“Blue” or “Red”) which instructed participants which color to attend to in that trial. The trial lasted for 3.25 seconds during which dots of either of the colors could move from 0 to 3 times in total. If the participants were instructed to attend to the blue dots and the blue dots moved coherently, they had to press the response button. In that case they would hear the auditory feedback signaling the correct detection of the motions. **B.** The experiment started with a practice and a baseline block in which the participants heard an audio cue at the beginning of the trial and two types of feedback sounds (incorrect or correct). In the training block a third sound was introduced to signal that the participants were both correct and received a reward for that response. They would still at times hear the old correct feedback which would signal that they were correct, but not rewarded. The test phase was the same as the baseline phase.

The experiment started with 4 practice blocks of 60 trials in each block. After each block, participants received feedback on their performance (percentage of correctly identified motions). During the practice blocks, participants performed the same task as in the main experiment (without rewards). After finishing the practice phase, participants completed 12 blocks (each consisting of 50 trials) divided into 3 phases (*baseline*, *training*, and *test*; *Figure 1B*) of 4 blocks each. Each phase contained 100 trials in which participants were instructed to attend to the red color and 100 trials in which they were instructed to attend to the blue color. Out of those 100 trials, 40 trials contained no dot motion, while 60 trials contained one, two, or three dot motions. The trials in which participants attended to one or the other color as well as the trials with different number of motions were randomly intermixed. Participants executed the coherent motion detection task, as described above, throughout all three phases (baseline, training, and test). In the training phase, participants could earn additional monetary rewards (up to 6 €) based on their actual performance. After completing the baseline phase, they were instructed that one of the colors would be paired with high probability (80%) and the other color with low probability (20%) of earning 10 extra cents for each correct motion detection. The mapping between color and reward probability was counterbalanced across participants. Receipt of the reward was signaled by a new tone that replaced the usual correct tone. If the correct tone was a sine wave of 800 Hz, the reward tone was a sine wave of 1,200 Hz (counterbalanced across participants). At the end of each of 4 training blocks, participants received feedback regarding both their performance and the amount of reward earned within the block (on average 5.5 € out of the maximal 6 € across all 4 blocks). The third phase (test) was identical to baseline and participants were explicitly informed that they would not be able to earn any more rewards. The entire task lasted for approximately 50 minutes, including short breaks in between blocks. Afterwards, participants completed two questionnaires aimed at assessing reward sensitivity (BIS-BAS; Franken et al., 2005) and depression levels (BDI-II; Van der Does, 2002). The collection of the questionnaire data is not reported here as it was collected for exploratory purposes in order to form a larger database of neural and self-report measures of reward processing. The experiment was implemented using Cogent Graphics developed by John Romaya at the LON at the Wellcome Department of Imaging Neuroscience.

### EEG recording and preprocessing

Electroencephalographic activity (EEG) was recorded with an ActiveTwo amplifier (BioSemi, Inc., The Netherlands) at a sampling rate of 512 Hz. Sixty-four Ag/AgCl electrodes were fitted into an elastic cap, following the international 10/10 system (Chatrian, Lettich, & Nelson, 1985). The common mode sense (CMS) active electrode and the driven right leg (DRL) passive electrode were used as reference and ground electrodes, respectively. Additional external electrodes were applied to the left and right mastoids, as well as on the outer canthi of each eye and in the inferior and superior areas of the left orbit (to record horizontal and vertical electrooculogram, EOG).

Data preprocessing was performed offline with custom MATLAB scripts and functions included in EEGLAB v14.1.1b (Delorme & Makeig, 2004). After subtracting the mean value of the signal (DC offset), the continuous EEG data were epoched between 0 and 3,250 ms, corresponding to the beginning and end of the trial, respectively. After referencing to *Cz*, FASTER v1.2.3b (Nolan, Whelan, & Reilly, 2010) was used for artifact identification and rejection using the following settings: (i) over the whole normalized EEG signal, channels with variance, mean correlation, and Hurst exponent exceeding *z* = ±3 were interpolated via a spherical spline procedure (Perrin, Pernier, Bertrand, & Echallier, 1989); (ii) the mean across channels was computed for each epoch and, if amplitude range, variance, and channel deviation exceeded *z* = ±3, the whole epoch was removed; (iii) within each epoch, channels with variance, median gradient, amplitude range, and channel deviation exceeding *z* = ±3 were interpolated; (iv) grand-averages with amplitude range, variance, channel deviation, and maximum EOG value exceeding *z* = ±3 were removed; (v) epochs containing more than 12 interpolated channels were discarded. Subsequently, automated routines were used to reject all trials with blinks or horizontal eye-movements exceeding 25 microvolts. For details, see our commented code at https://osf.io/kjds3/. After preprocessing, the average number of interpolated channels was 3.61 (*SD* = 1.23, range 1 – 6) and the mean percentage of rejected epochs was 8.77% (*SD* = 6.71, range 0 – 27.78). After re-referencing to averaged mastoids, trials in each condition were averaged separately for each participant, resulting in the following conditions: (i) baseline, red attended; (ii) baseline, blue attended; (iii) training, red attended; (iv) training, blue attended; (v) test, red attended; (vi) test, blue attended.

After removing linear trends, SSVEP amplitudes were computed as the absolute of the complex Fourier coefficients of the trial-averaged EEG in a time-window from 500 ms (to exclude the typically strong phasic visual evoked response to picture onset) to 3,250 ms after stimulus onset. Electrodes with maximum SSVEP amplitudes were identified by calculating isocontour voltage maps based on grand-averaged data collapsed across all conditions. This procedure identified a cluster consisting of the four electrodes Oz, O2, POz, and Iz, which were chosen for further analysis. SSVEP amplitudes were normalized (rescaled) for each participant and frequency (10 and 12 Hz) separately by dividing amplitudes by the average amplitude of the two conditions in the baseline.

### Statistical analyses

Behavioral and EEG data were analyzed using Bayesian multilevel regressions. We fitted and compared multiple models of varying complexity to predict observer sensitivity, reaction times for correct responses, and SSVEP amplitudes. For the behavioral data, mean reaction times of correct detections (hits) and sensitivity (d′) were analyzed. Sensitivity index d′ (Macmillan & Creelman, 2004) was calculated with adjustments for extreme values (Hautus, 1995) using the *psycho* R package (for the method see: Pallier, 2002). When calculating d′, responses to the coherent motion of the attended color were considered as hits, while responses to the coherent motion of the unattended color were considered as false alarms.

Each fitted model included both constant and varying effects (also known as fixed and random). Participant-specific characteristics are known to affect both behavioral performance (e.g., response speed) and EEG signal (e.g., skull thickness, skin conductance, hair); therefore, we accounted for this variability by adding varying intercepts in our models. Additionally, the studied effects (i.e., selective attention and reward sensitivity) are known to vary in magnitude over participants, so we opted for including varying slopes in our models^2^.

Models were fitted in R using the *brms* package (Bürkner, 2016) which employs the probabilistic programming language *Stan* (Carpenter et al., 2016) to implement Markov Chain Monte Carlo (MCMC) algorithms in order to estimate posterior distributions of the parameters of interest (details about the fitted models can be found in the data analysis scripts at https://osf.io/kjds3/). Each model was fitted using weakly informative prior distributions (described below) and Gaussian likelihood. Four MCMC simulations (“chains”) with 6,000 iterations (3,000 warmup) and no thinning were run to estimate parameters in each of the fitted models. Further analyses were done following the recommendations for Bayesian multilevel modeling using *brms* (Bürkner, 2016, 2017; Nalborczyk & Bürkner, 2019). We confirmed that all models converged by examining trace plots, autocorrelation, and variance between chains (Gelman-Rubin statistic; Gelman & Rubin, 1992). We compared models based on their fit to the actual data using the Bayesian *R*^2^ (Gelman, Goodrich, Gabry, & Ali, 2017), and their out-of-sample predictive performance using the Widely Applicable Information Criterion (WAIC; Watanabe, 2010). The best model was selected and the posterior distributions of conditions of interest were examined. Differences between conditions were assessed by computing the mean and the 95% highest density interval (HDI) of the difference between posterior distributions of the respective conditions (Kruschke, 2014). Additionally, we calculated the evidence ratios (ERs) for our hypotheses as the ratios between the percentage of posterior samples on each side of the zero of the difference distribution between two conditions. ERs represent the ratio between the probability of a hypothesis (e.g. “Condition A is larger than condition B”) against its alternative (“Condition B is larger than condition A”). As a rule of thumb, we interpreted our results as providing “inconclusive” evidence when 1 < ER < 3, “anecdotal” evidence when 3 < ER < 10, and “strong” evidence when ER > 10. When ER > 12000 (the maximum number of posterior samples), the posterior distribution was completely on one side of zero, thus providing “very strong” evidence.

#### Behavioral data

We fitted three models to predict sensitivity (d′) and reaction times (in milliseconds) separately (see *Figure 2* for the raw data and *Supplementary Table 1* for the descriptive statistics). First, we fitted the *Null model* with a constant and varying intercepts across participants. This model was fitted in order to explore the possibility that the data would be best explained by simple random variation between participants. To investigate the effect of reward phase (baseline, training, test), we fitted the *Reward phase* model which included only reward phase as the constant predictor, as well as varying intercepts and slopes across participants for this effect. To investigate the possible interaction between reward phase and reward, we fitted the *Reward phase × Reward Probability* model including the intercepts and slopes of these two effects and their interaction as both constant and varying effects. All models had a Gaussian distribution as the prior for the intercept (for sensitivity: centered at 1.8 with a standard deviation of 1; for reaction times: centered at 500 with a standard deviation of 200). The models with slopes also included a Gaussian distribution as prior for the slopes (for sensitivity: centered at 0 with a standard deviation of 2; for reaction times: centered at 0 with a standard deviation of 200). The means for the priors for the intercepts were selected based on a previous study with a similar task (Andersen & Müller, 2010). The standard deviations of all of the prior distributions were chosen so that the distributions are very wide and thus only weakly informative. Note that there are two additional models that, although possible to fit, are not plausible in the context of our experiment. Specifically, the model including only the effect of reward probability overlooks the fact that this effect would necessarily be most pronounced in the training phase, thus interacting with the effect of reward phase. The same logic applies to the model with additive effects of reward phase and probability (i.e., these effects could not act independently in our experimental design).

**Figure 2.**
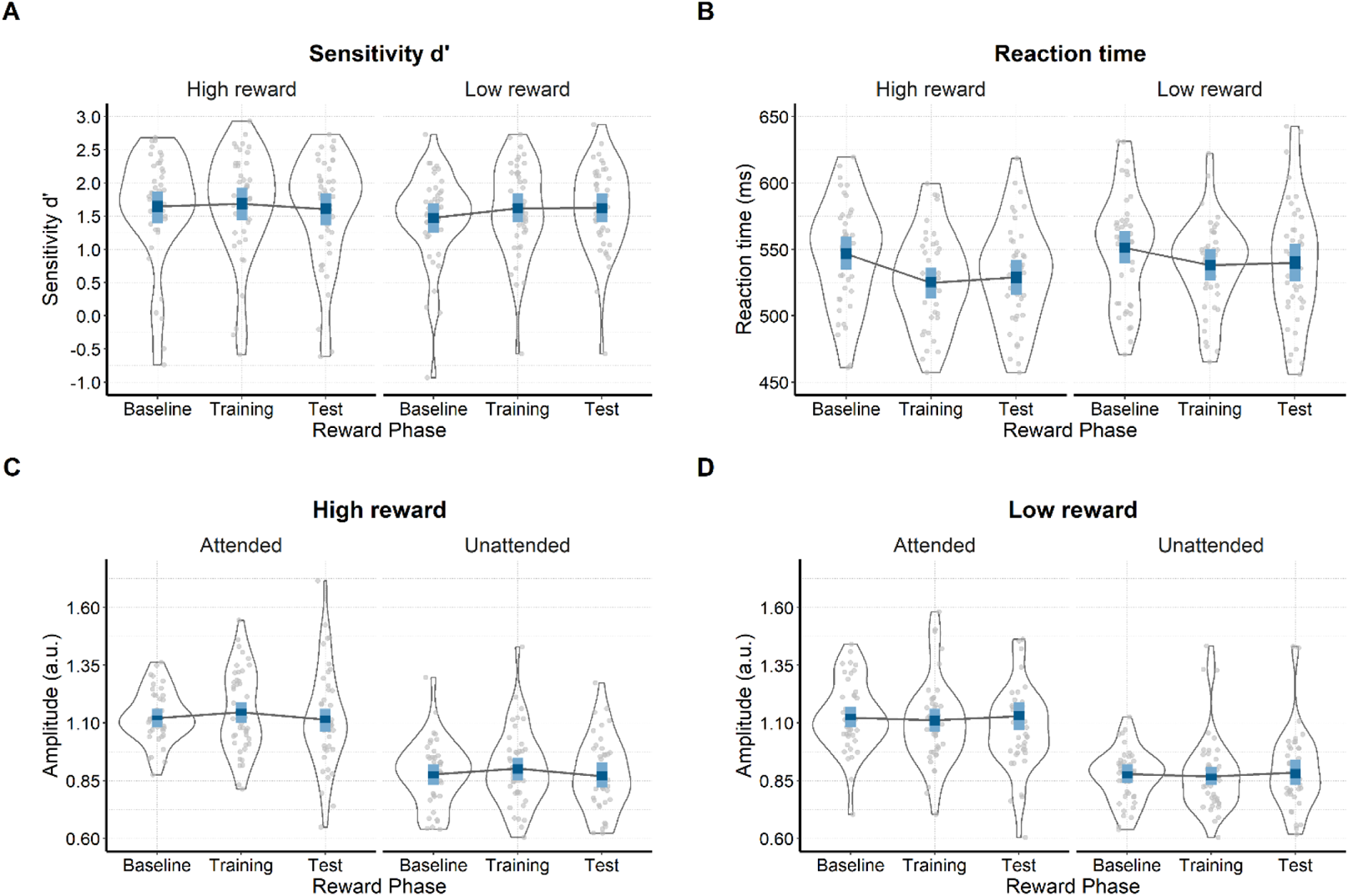
Raw and modelled data. Violin plots displaying raw data for each participant (grey dots), separately for each condition. Results from the winning models are presented in blue (dark blue – 50% HDIs and light blue – 95% HDIs). **A.** Sensitivity (d′) **B.** Reaction times (ms) **C.** SSVEP amplitudes (arbitrary units) in response to the color related to high reward on trials in which it is attended or unattended. **D.** SSVEP amplitudes for the color linked to low reward on trials when it was attended or unattended.

#### SSVEP amplitudes

We fitted seven models to predict the trial-averaged SSVEP amplitudes (in a.u. due to the normalization) across conditions (see *Figure 2C, Figure 2D*, and *Supplementary Table 2*). The *Null model* included one constant and varying intercepts across participants. The *Attention model* included attention as predictor; the *Reward Phase model* included the effect of reward phase; the *Reward Phase + Attention* model included the additive effects of reward phase and attention; and the *Reward Phase × Attention* model also included the interaction between reward phase and attention. The *Reward probability × Reward phase + Attention* model consisted of the effects of reward and phase, their interaction, and the independent effect of attention. The last model was the *Reward probability × Reward phase × Attention* model which included all predictors and their interaction. All models, except for the *Null model*, included varying intercepts and slopes across participants for all effects. All models included a Gaussian distribution as the prior for the intercept (centered at 1 with a standard deviation of 1). The mean across both attended and unattended conditions is approximately 1 in this paradigm (Andersen & Müller, 2010), while the normalized amplitudes are in the 0-2 range (the normalized amplitude of 2 for the attended stimulus would equal the physical removal of the unattended stimulus), which is why we opted for the standard deviation of 1 for the prior distributions. In addition, the models with slopes included a Gaussian distribution as the prior for the slopes (centered at 0 with a standard deviation of 1). As was the case for the behavioral data, several models were not fitted because they were not plausible in the context of our experiment (i.e., models that include both reward phase and probability, but not their interaction, are implausible because reward probability could not affect the baseline phase as the reward mapping information was provided upon completion of the baseline).

## Results

### Behavioral results

#### Sensitivity d′

The analyses of sensitivity revealed that participants successfully performed the task, as d′ was well above chance level across all conditions. Of all the tested models, the *Reward phase × Reward probability* model best predicted sensitivity (*Table 1*). The posterior distributions of the interaction model (*Figure 2A* and *Table 2*) revealed that sensitivity improved in the training phase compared to the baseline for low reward (*M* = 0.14; 95% HDI [0.01, 0.27]; ER = 57.82), while the improvement for the high reward color was in the same direction, but not statistically robust (*M* = 0.04; 95% HDI [−0.08, 0.17]; ER = 3.10). This improvement was slightly more pronounced for low compared to high reward (*M* = 0.10; 95% HDI [−0.08, 0.27]; ER = 6.25). Conversely, there was no evidence for a difference between training and test phases in the low reward condition (*M* = 0.00; 95% HDI [−0.13, 0.13]; ER = 1.09), while there was a reduction in sensitivity in the high reward condition (*M* = −0.08; 95% HDI [−0.20, 0.05]; ER = 8.52). These results suggest higher sensitivity for coherent motion detection in the training phase compared to baseline, which was more pronounced for the low relative to the high reward color. This somewhat counterintuitive effect could be explained by the faster reaction times to the high compared to the low reward color, which we focus on in the following section. Finally, we found very little evidence of a change in sensitivity from the training to the test phase. Importantly we found a baseline difference between the high and low reward conditions (*Table 2*). This result is likely due to random fluctuations because in the baseline phase participants are not aware of any reward contingencies. While this result does not affect our interpretation because we analyze the change in each of the two colors separately across the phases of the experiment, the magnitude of the baseline difference suggests that the effects of reward on sensitivity are rather small. This is in line with previous work on value-driven attention in which the reward-driven effects are more commonly reflected in reaction times rather than changes in accuracy (Anderson, 2016; Awh et al., 2012; Chelazzi et al., 2013; Failing & Theeuwes, 2017).

**Table 1.**
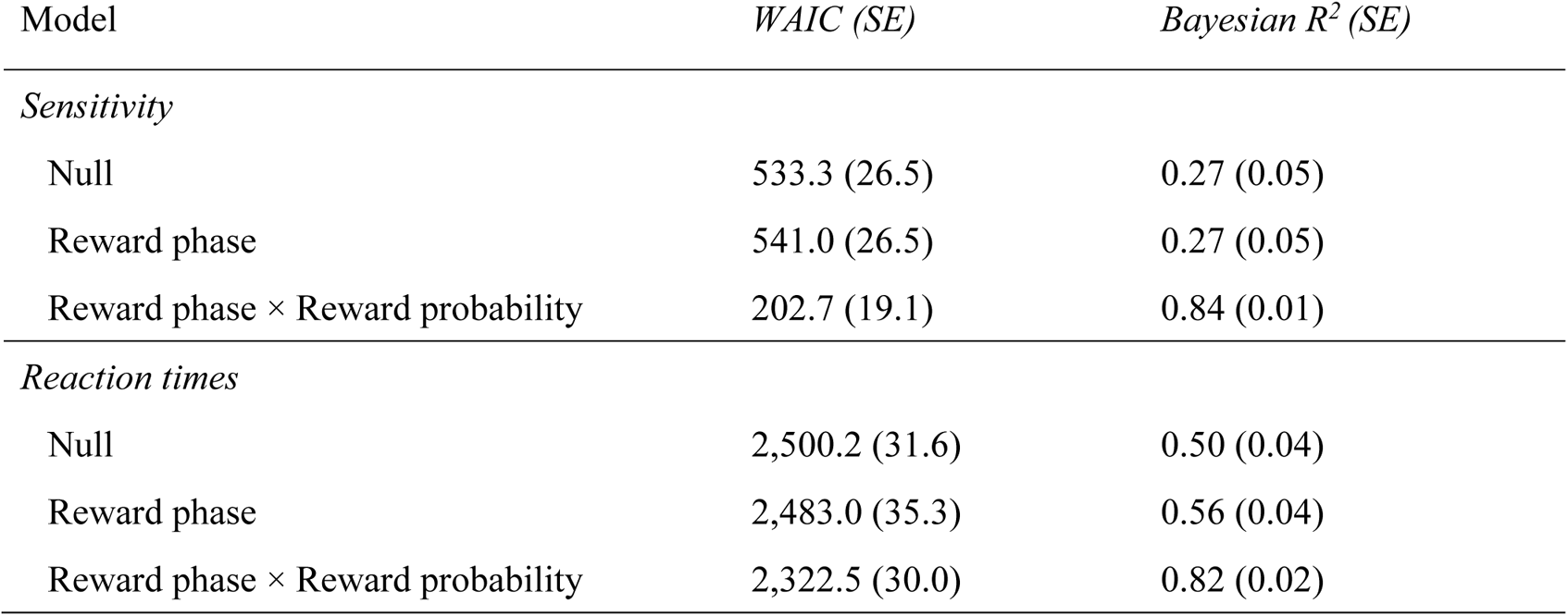
Mean and standard errors (in parenthesis) of WAIC and Bayesian R^2^ for each model predicting sensitivity and reaction times.

**Table 2.** Means and 95% HDIs of the posterior distributions of reaction times and sensitivity in each condition.

| Reward phase | Reward probability | Sensitivity ($d'$) | Reaction times (milliseconds) |
| --- | --- | --- | --- |
| Baseline | High | 1.64 [1.39, 1.87] | 546.54 [534.33, 559.30] |
| Baseline | Low | 1.48 [1.25, 1.69] | 551.13 [539.34, 563.50] |
| Training | High | 1.69 [1.44, 1.93] | 524.91 [512.94, 536.30] |
| Training | Low | 1.62 [1.41, 1.84] | 537.99 [526.48, 550.32] |
| Test | High | 1.61 [1.36, 1.84] | 528.97 [515.90, 541.99] |
| Test | Low | 1.62 [1.41, 1.84] | 539.85 [525.63, 554.34] |

#### Reaction times

The *Reward phase × Reward probability* model best predicted reaction times (*Figure 2B* and *Table 1*). In the training, compared to the baseline phase, participants were reliably faster in detecting the motions of both the high (*M* = −21.60 ms; 95% HDI [−29.90, −12.80]; ER > 12,000, i.e., the whole posterior distribution was below zero thus the ER is larger than the total number of posterior samples) and the low reward colors (*M* = −13.10 ms; 95% HDI [−21.70, −4.69]; ER = 999). Moreover, this difference between baseline and training was larger for detecting motions of high relative to low reward color (*M* = −8.49 ms; 95% HDI [−18.60, 2.06]; ER = 17.18). We found weak evidence for changes in reaction times between the training and the test phase. There was a very small, but not statistically robust, increase in reaction times in the test compared to training phase for the high reward color (*M* = 4.07 ms; 95% HDI [−4.52, 13.10]; ER = 4.40), and no difference for the low reward color (*M* = 1.87 ms; 95% HDI [−6.93, 10.70]; ER = 1.98). We confirmed that the reward-induced changes persisted even after rewards were no longer available by comparing the reaction times in the baseline phase to the test phase. These analyses revealed that participants responded faster in the test phase relative to the baseline phase to both high (*M* = −17.60 ms; 95% HDI [−28.40, −6.23]; ER = 999) and low reward stimuli (*M* = −11.30 ms; 95% HDI [−22.60, −0.72]; ER = 44.45). Further, this speeding up was more pronounced for the stimuli previously related to high compared to low reward probability (*M* = −6.29 ms; 95% HDI [−16.30, 4.44]; ER = 7.70). These results indicate that participants were faster in detecting coherent motions in the condition in which they could earn rewards (training), and more so for high than low reward color. Also, there was a small increase in reaction times for the high reward condition and no difference in the low reward condition when the rewards were no longer available (test). Crucially, this increase was limited, and participants were still faster to respond in the test compared to the baseline phase, and more so for the stimuli related to high compared to low reward probability. Supplementary analyses carried out to assess possible training effects indicated some evidence for the presence of training effects in sensitivity and scant evidence for such effects in reaction times (*Supplementary materials*).

### SSVEP amplitudes

As shown in *Figure 3*, SSVEP amplitudes averaged over conditions peaked at central occipital channels (i.e., *Oz*, *POz*, *O2*, *Iz*). Also, the amplitude spectra showed the expected pronounced peaks at the frequencies of 10 and 12 Hz.

**Figure 3.**
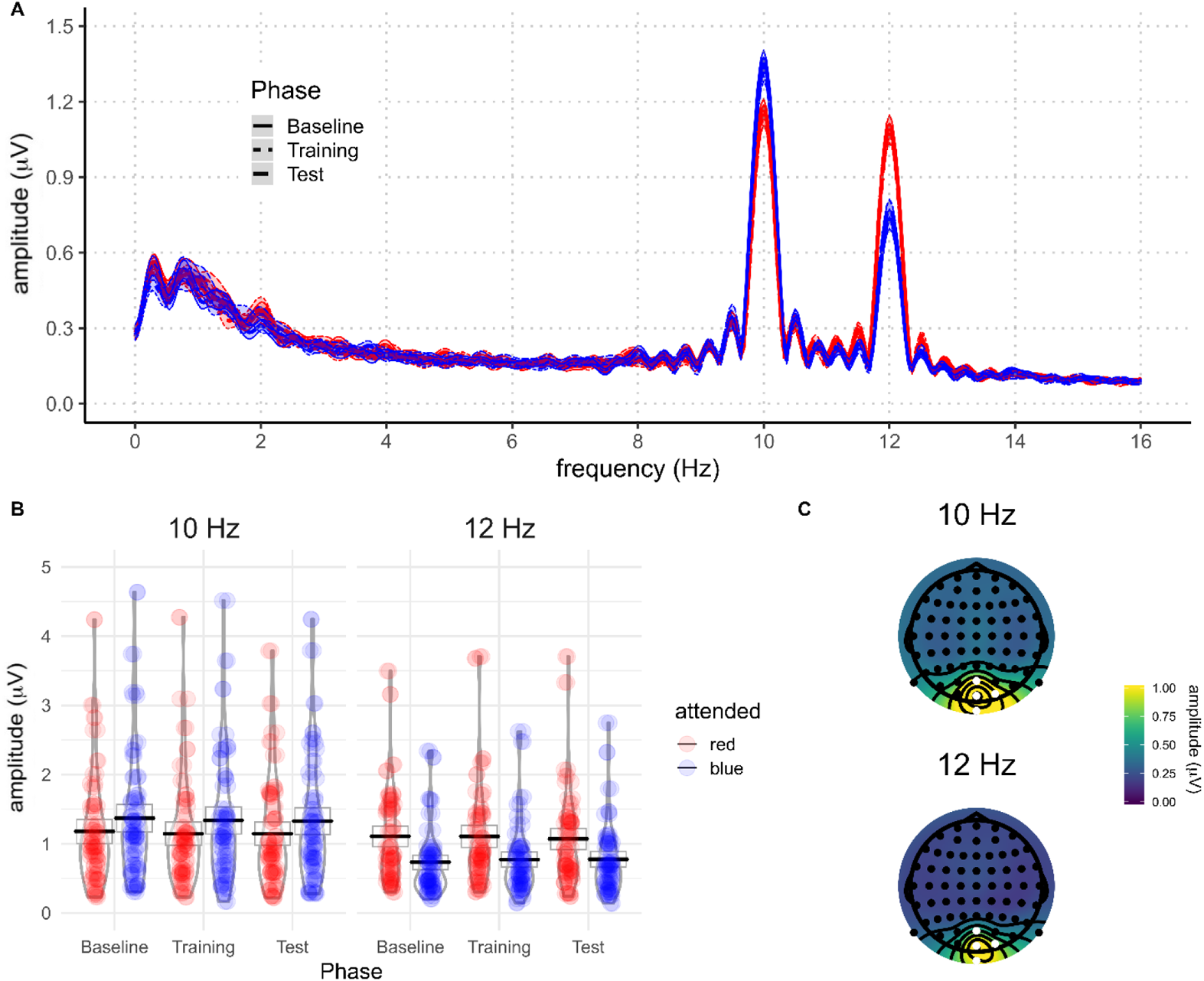
A) Grand average amplitude spectra (only for visualization purposes, 1 Hz high-pass FIR filter and zero-padded to 8 times the length of the data) derived from EEG signals at best four-electrode cluster plotted for the different experimental conditions (blue: attended; red: unattended; solid: baseline phase; dotted: rewarded phase; dashed: non-rewarded phase). The shaded areas around the means indicate 95% confidence intervals. **B)** Individual and average amplitudes (with 95% confidence intervals) for blue (10 Hz) and red (12 Hz) across task conditions. **C)** Topographies of SSVEP amplitudes, averaged across all participants and conditions, at 10 Hz and 12 Hz. Electrodes selected for the analysis are highlighted in white.

The *Reward probability × Reward phase + Attention* model best predicted SSVEP amplitudes across conditions (*Table 3*). However, the *Reward probability × Reward phase × Attention* had only slightly lower explanatory power relative to the winning model. Here we draw inferences from the winning model, but note that the conclusions do not substantially change when analyzing the model which includes the three-way interaction. The analysis of the posterior distributions of the winning model (*Figure 2* and *Table 3*) revealed a very strong effect of voluntary selective attention, indicating that participants were following the instructions and attending the dots of the cued color. Across all conditions, SSVEP amplitudes were higher when the eliciting stimulus was attended compared to when it was unattended. In the winning model, this effect did not interact with the other factors in the model, i.e., the magnitude of selective attention was unaffected by reward probability and reward phase. The posterior distribution of the difference between attended and unattended stimuli did not include zero, revealing a very strong effect of voluntary attention. Namely, the attended stimuli very reliably elicited higher SSVEP amplitudes compared to the unattended ones (*M* = 0.24; 95% HDI [0.20, 0.29]; ER > 12,000). These results reveal a very robust effect of voluntary selective attention across all experimental conditions: the SSVEP response was systematically larger when the driving stimulus was attended.

**Table 3.** Model comparison indices for EEG results

| Model | *WAIC (SE)* | *Bayesian R<sup>2</sup> (SE)* |
| --- | --- | --- |
| Null | -22.3 (56.2) | 0.01 (0.01) |
| Reward phase | -31.8 (55.0) | 0.05 (0.01) |
| Attention | -436.5 (66.4) | 0.37 (0.02) |
| Reward phase + Attention | -464.7 (64.9) | 0.40 (0.02) |
| Reward phase × Attention | -461.3 (65.2) | 0.41 (0.02) |
| Reward probability × Reward phase + Attention | -696.1 (71.9) | 0.55 (0.02) |
| Reward probability × Reward phase × Attention | -690.4 (71.9) | 0.55 (0.02) |

**Table 4.** Means and 95% HDIs of the posterior distributions of the SSVEP amplitudes for each condition.

| Attention | Reward phase | Reward probability | Amplitudes (a.u.) |
| --- | --- | --- | --- |
| Attended | Baseline | High | 1.12 [1.08, 1.16] |
| Attended | Baseline | Low | 1.12 [1.07, 1.17] |
| Attended | Training | High | 1.15 [1.10, 1.19] |
| Attended | Training | Low | 1.11 [1.07, 1.16] |
| Attended | Test | High | 1.11 [1.06, 1.17] |
| Attended | Test | Low | 1.13 [1.07, 1.19] |
| Unattended | Baseline | High | 0.88 [0.83, 0.92] |
| Unattended | Baseline | Low | 0.88 [0.84, 0.92] |
| Unattended | Training | High | 0.90 [0.85, 0.95] |
| Unattended | Training | Low | 0.87 [0.82, 0.91] |
| Unattended | Test | High | 0.87 [0.82, 0.92] |
| Unattended | Test | Low | 0.88 [0.83, 0.94] |

The winning model also included the interaction between reward phase and reward probability, but this interaction remained the same for both attended and unattended stimuli. SSVEP amplitudes were higher in the training phase than at baseline for the high reward color (*M* = 0.02; 95% HDI [−0.01, 0.06]; ER = 9.53), both when it was attended and unattended. However, there was no evidence of difference for the change in SSVEP amplitudes from baseline to training for the low reward color (*M* = 0.01; 95% HDI [−0.03, 0.05]; ER = 2.58). Comparing the training to the test phase, the amplitudes of the high reward color were reduced (*M* = −0.03; 95% HDI [−0.07, 0.01]; ER = 13.71), while the amplitudes of the low reward color did not substantially change (*M* = −0.02; 95% HDI [−0.06, 0.02]; ER = 3.72).

To summarize, visual processing of the high reward color stimulus was enhanced in the phase in which the participants could earn monetary rewards. This gain in neural processing returned to baseline in the subsequent test phase in which the rewards were no longer available. Importantly, the reward-dependent modulation of the visual cortex activity occurred irrespective of whether that color was attended or not, i.e., it did not affect voluntary allocation of attention to the cued color. Finally, visual processing of the low reward color remained constant across the three phases of the experiment.

## Discussion

In this study we investigated the neural mechanisms through which voluntary selective attention and reward history jointly guide visual processing. We compared the processing of attended and unattended stimuli of different reward probabilities on a continuous global motion discrimination task. Compared to baseline, the introduction of rewards sped up task performance, especially for the higher reward stimuli, which was accompanied by enhanced processing of these stimuli in the visual cortex (as suggested by higher SSVEP amplitude values). This sensory gain was present both when the high reward stimulus was attended and unattended, thus indicating that rewards influenced visual processing independently of voluntary selective attention. When rewards were no longer available, sensory processing of high reward stimuli returned to baseline levels, but participants were still faster to detect coherent motion of high vs. low reward stimuli relative to the baseline.

The introduction of rewards improved behavioral performance on the task and facilitated the visual processing of stimuli associated with high rewards. This effect on SSVEP amplitudes is likely localized in the V1-V3 areas of the visual cortex, as reported in previous studies using the same task that conducted formal source analysis of the SSVEP (Andersen et al., 2009; Andersen & Müller, 2010; Andersen, Hillyard, & Müller, 2008). This effect was the same both when the high reward stimulus was attended and unattended. Thus, this effect was independent of the effect of voluntary selective attention as reflected in the enhanced processing of the attended compared to unattended stimuli (Andersen & Müller, 2010). This pattern of results suggests that the effect of reward acted independently of voluntary attention, which is in line with previous work showing the independent influence of reward and task-relevance on stimulus processing in the extrastriate visual cortex (Buschschulte et al., 2014; Garcia-Lazaro et al., 2019). This finding supports the predictions of the models which propose that the effect of reward history on visual processing is independent from voluntary attention (Anderson, 2016; Awh et al., 2012; Chelazzi et al., 2013; Failing & Theeuwes, 2017). Further, this finding can help refine models highlighting the role of rewards in the allocation of cognitive control. These models (Brown & Alexander, 2017; Holroyd & McClure, 2015; Shenhav et al., 2013; Verguts et al., 2015) are largely focused on activity in the frontoparietal regions, for example the dorsolateral prefrontal cortex and the anterior cingulate cortex, which are known to increase their activation in anticipation of rewards (Krebs, Boehler, Roberts, Song, & Woldorff, 2012; Pessoa & Engelmann, 2010; Schevernels, Krebs, Santens, Woldorff, & Boehler, 2014). However, these models are not explicit about their predictions of how top-down signals from these areas modulate the processing of stimuli at the level of the visual cortex. Our findings suggest that increased rewards act to enhance the processing of the stimuli related to high rewards independently of other top-down voluntary attention effects, which is similar to the way in which physical salience of stimuli (i.e., contrast) acts in the same paradigm (Andersen et al., 2012). Interestingly, this is at odds with recent finding showing that a flagship cognitive control effect, post-error adjustments, operates through enhancement of voluntary selective attention as measured by SSVEPs using an adapted version of the task used here (Steinhauser & Andersen, 2019). This indicates a possible dissociation between the effects of reward and other cognitive control effects on selective attention. Dissociations between cognitive control and reward effects should be further addressed, both theoretically and empirically.

In the test phase, behavioral performance displayed similar patterns as in the training phase. Individuals were faster to detect motions of the dots in color related to high compared to low reward. This finding follows the reward-history effects reported in several paradigms (Anderson, Laurent, & Yantis, 2011; Della Libera & Chelazzi, 2009; Failing & Theeuwes, 2014). However, our SSVEP results show that the visual processing of high reward stimuli returned to baseline levels, diverging from the behavioral pattern of results. This may indicate that the longer lasting effect of reward history was not mediated by the prolonged gain enhancement in sensory processing as measured by the SSVEPs, contrary to the predictions of the models accounting for the effects of reward history on attention (Anderson, 2016; Awh et al., 2012; Chelazzi et al., 2013; Failing & Theeuwes, 2017). This result is predicted by models which relate cognitive control and reward, as they predict that reward-related enhancements should return to baseline levels when rewards are no longer available (Brown & Alexander, 2017; Holroyd & McClure, 2015; Shenhav et al., 2013; Verguts et al., 2015). This finding suggest that visual processing can be adapted in a much more flexible way than predicted by the models focused on the reward-history effects on attention. Of note, it is possible that our SSVEP measure captures more sustained processing of features in visual cortex, while the effects of reward history could be specifically locked to the onset of the rewarded stimulus (Donohue et al., 2016; Hickey et al., 2010; Luque et al., 2017; MacLean & Giesbrecht, 2015). However, there are at least two studies which have not found evidence for the effects of reward history on early visual processing (Qi et al., 2013; Tankelevitch et al., 2020). This leaves open the possibility that effects of reward history are not necessarily driven purely by gains in sensory processing. One interesting possibility, which should be explored in future studies, is that rewards initially improve performance by enhancing stimulus salience, but later rely on more direct stimulus-response mappings. Finally, it is important to note that our paradigm involves a cue on every trial which induces a direct goal, at odds with most studies assessing the influence of reward-history on attention. Further research using SSVEPs ought to be conducted in order to explicitly address effects of reward history on SSVEP amplitudes.

Our paradigm allowed us to simultaneously measure the processing of stimuli linked to both high and low value. Some initial evidence for attentional suppression of stimuli linked to low compared to high rewards has been found at the behavioral and neural level (Hickey & Peelen, 2015; Padmala & Pessoa, 2011). Suppression of visual features linked to low or no rewards has also been proposed as one of the potential mechanisms through which incentives impact attention (Chelazzi et al., 2013; Anderson, 2016; Failing & Theeuwes, 2018). On the contrary, in this study we found no evidence for this proposal. Suppression was neither observed when the low value color was attended, nor when it was unattended. Visual processing of the low reward color, as indexed by SSVEP amplitudes, was strongly affected by attention, but remained unchanged by reward throughout the experiment. There are three features of our experiment that may explain this finding. First, in our experiment both colors were related to rewards, but they differed in reward value. Conversely, Hickey and Peelen (2015) showed evidence for the suppression of the non-rewarded feature for objects which were never rewarded. In our paradigm, it could be beneficial for participants not to suppress the low value color because correct responses to the motions of this color would still earn them a reward on 20% of trials. Second, in our experiment the attended color changed on a trial-by-trial basis, whereas the experiment of Hickey and Peelen (2015) consisted out of small blocks of 16 trials in which the attended object was always the same (e.g., searching for a car in a complex picture). When searching for one object or feature across a number of future trials, it is possible that the optimal solution for the visual system is to suppress the processing of the other features or objects (i.e., goal-irrelevant stimuli). However, if the attended feature is likely to change on each trial, as in our experiment, the suppression of the low rewarded feature could be maladaptive as it would carry a cost of reconfiguring the control signals on every trial (for a computational implementation of a reconfiguration cost see: Musslick, Shenhav, Botvinick, & Cohen, 2015). Third, our experiment included a shorter training phase compared to some of the previous experiments which demonstrated reliable behavioral effects of the value-driven attentional bias (Anderson, Laurent, & Yantis, 2011; Della Libera & Chelazzi, 2009; Failing & Theeuwes, 2014). While the lower number of reward-stimulus pairings (120 for high and low reward colors each here) could lead to weaker effects, we were still able to conceptually replicate the previous behavioral findings, indicating that we were successful at inducing a reward-driven bias. However, we cannot fully exclude the possibility that sustained effect of rewards at the neural level would have been observed with a longer training phase.

The design of this study and the use of the SSVEPs allowed us to independently assess the influence of voluntary attention and reward on sensory processing in the visual cortex. This enabled us to directly compare the magnitude of these two factors on sensory processing. While both modulated visual processing, it is important to note that the effect of voluntary attention on visual processing (30% increase for the attended vs. the unattended stimuli; based on the regression weights from the fitted models) was an order of magnitude stronger than the effect of reward (3% increase from baseline to training for the high reward stimuli). Thus, even though reward associations can influence processing in opposition to voluntary attention, our results suggest that the magnitude of this effect is very small compared to the effect of voluntary attention. Most theoretical models to date have focused on how top-down and reward-driven attention jointly guide stimulus processing (Awh et al., 2012), but how much each of these processes contribute to stimulus processing still has to be incorporated into these theoretical models. This finding is especially important in the light of recent studies investigating the relevance of reward-driven automatic biases in attention in clinical disorders such as addiction (Anderson, 2016) and depression (Anderson, Leal, Hall, Yassa, & Yantis, 2014). While it is possible that more automatic biases in attention play a role in these disorders, it is also important to focus on the influence of more goal-directed processes which are likely to have a bigger impact on cognition in clinical disorders (Grahek, Shenhav, Musslick, Krebs, & Koster, 2019).

In conclusion, in this study we directly assessed how voluntary attention and reward jointly guide attention. Our findings provide a novel insight into the flexible dynamics of visual processing by demonstrating that rewards can act independently of voluntary attention to enhance sensory processing in the visual cortex. However, sensory processing is flexibly readjusted when rewards are no longer available. This result suggests that top-down and reward effects independently affect sensory gain in the visual cortex which needs to be accounted for in theoretical models of motivation-cognition interactions. The effect can be flexibly removed as soon as the reward structure in the environment changes.

## Supplementary materials

### Means of the raw behavioral and SSVEP data

**Supplementary Table 1.** Means and 95% HDIs (in square brackets) of the raw data for sensitivity and reaction times

| Reward phase | Value | Sensitivity (d') | Reaction times (milliseconds) |
| --- | --- | --- | --- |
| Baseline | High | 1.64 [-0.04, 2.68] | 546.59 [485.64, 619.34] |
| Baseline | Low | 1.47 [ 0.04, 2.30] | 551.10 [490.50, 631.36] |
| Training | High | 1.69 [ 0.29, 2.73] | 524.99 [467.12, 599.49] |
| Training | Low | 1.62 [ 0.46, 2.68] | 537.94 [465.32, 584.63] |
| Test | High | 1.60 [-0.20, 2.73] | 528.98 [457.08, 599.83] |
| Test | Low | 1.62 [ 0.74, 2.88] | 539.75 [455.80, 623.21] |

**Supplementary Table 2.** Means and 95% HDIs of the raw data for the recorded SSVEP amplitudes in each condition

| Attention | Reward phase | Value | Amplitudes (a.u.) |
| --- | --- | --- | --- |
| Attended | Baseline | High | 1.13 [0.92, 1.52] |
| Attended | Baseline | Low | 1.13 [0.86, 1.52] |
| Attended | Training | High | 1.16 [0.80, 1.60] |
| Attended | Training | Low | 1.13 [0.76, 1.71] |
| Attended | Test | High | 1.13 [0.61, 1.61] |
| Attended | Test | Low | 1.13 [0.59, 1.84] |
| Unattended | Baseline | High | 0.87 [0.47, 1.17] |
| Unattended | Baseline | Low | 0.87 [0.49, 1.11] |
| Unattended | Training | High | 0.91 [0.54, 1.38] |
| Unattended | Training | Low | 0.89 [0.50, 1.28] |
| Unattended | Test | High | 0.88 [0.48, 1.23] |
| Unattended | Test | Low | 0.91 [0.44, 1.42] |

### Additional analyses to assess possible training effects

In order to assess potential training effects on behavioral performance, we split each reward phase into two halves (*Supplementary Figure 1* and *Supplementary Table 3*). If training effects were influencing the behavioral outcome, we could expect performance improvement through baseline and training. To investigate this possibility, we fitted the *Reward phase × Value* model that was identical to the one described in the results section. We then compared behavioral performance between the first and the second part of the baseline phase, and between the second part of baseline and the first part of training phase.

**Supplementary Table 3.** Means and 95% HDIs of the raw data for sensitivity and reaction times across six phases of the experiment

| Reward phase | Value | Sensitivity (d') | Reaction times (milliseconds) |
| --- | --- | --- | --- |
| Baseline1 | High | 1.48 [-0.36, 2.62] | 548.84 [479.43, 613.76] |
| Baseline1 | Low | 1.32 [ 0.09, 2.35] | 548.43 [458.26, 610.63] |
| Baseline2 | High | 1.60 [-0.27, 2.56] | 544.34 [454.56, 620.36] |
| Baseline2 | Low | 1.47 [ 0.08, 2.33] | 554.01 [479.48, 632.80] |
| Training1 | High | 1.54 [-0.08, 2.65] | 521.40 [437.90, 587.57] |
| Training1 | Low | 1.59 [ 0.47, 2.45] | 542.34 [463.65, 593.47] |
| Training2 | High | 1.59 [ 0.08, 2.56] | 528.74 [462.00, 598.58] |
| Training2 | Low | 1.48 [ 0.00, 2.62] | 533.94 [479.38, 618.25] |
| Test1 | High | 1.48 [-0.07, 2.47] | 528.58 [457.88, 596.17] |
| Test1 | Low | 1.50 [ 0.36, 2.50] | 536.54 [444.89, 621.00] |
| Test2 | High | 1.49 [-0.38, 2.49] | 529.30 [448.24, 606.00] |
| Test2 | Low | 1.55 [ 0.65, 2.55] | 543.01 [450.11, 617.44] |

The posterior distributions for sensitivity (*Supplementary Figure 1A* and *Supplementary Table 4*) revealed performance improvement from the first to the second part of the baseline for both high (*M* = 0.12; 95% HDI [−0.05, 0.28]; ER = 11.05) and low (*M* = 0.15; 95% HDI [0.01, 0.32]; ER = 36.04) value conditions. When comparing the second part of baseline to the first part of training, there was only a very small improvement in sensitivity in the high value condition (*M* = 0.06; 95% HDI [−0.11, 0.22]; ER = 2.94), and a much bigger one in the low value condition (*M* = 0.11; 95% HDI [−0.04, 0.28]; ER = 10.90). These results indicate that participants improved not only throughout the baseline phase, but also from the end of baseline to the first part of the training (albeit for low rewarded color only). This might indicate some presence of training effects in the sensitivity data.

The posterior distributions of reaction times (*Supplementary Figure 1B* and *Supplementary Table 2*) revealed only a very small difference between the first and the second part of baseline for high (*M* = −4.52; 95% HDI [−15.0,0 5.77]; ER = 4.21) value condition, while the low value condition was slightly slower in the second part of the baseline (*M* = 5.60 95% HDI [−4.76, 16.20]; ER = 5.71). The comparison between the second part of baseline and the first part of training revealed a very reliable speeding in both high (*M* = 22.90; 95% HDI [12.60, 33.80]; ER > 6000) and low (*M* = 11.60; 95% HDI [0.70, 22.10]; ER = 57.82) value conditions. These results clearly point to the absence of large training effects in the reaction time data.

Taken together, these results indicate that our effects were not driven by the improved performance over the course of the task. Although there is some evidence that sensitivity was improving during the baseline phase, reaction times clearly indicate that the main shift in performance happens in the beginning of training, when rewards are introduced. Importantly, the strongest behavioral effects in our study were found on reaction time data, as indicated in the *Results* section of the main text.

Similar analyses could not be performed for the EEG data, because splitting the number of trials in each phase would significantly affect the signal-to-noise ratio. However, our EEG results point to changes in SSVEP amplitudes in only one of the value conditions. If amplitude changes were mainly driven by training effects, the differences across reward phases would be expected for both value conditions. This observation, combined with the lack of strong training effects in behavior, suggests that our EEG results are not driven by training effects.

**Supplementary Figure 1.**
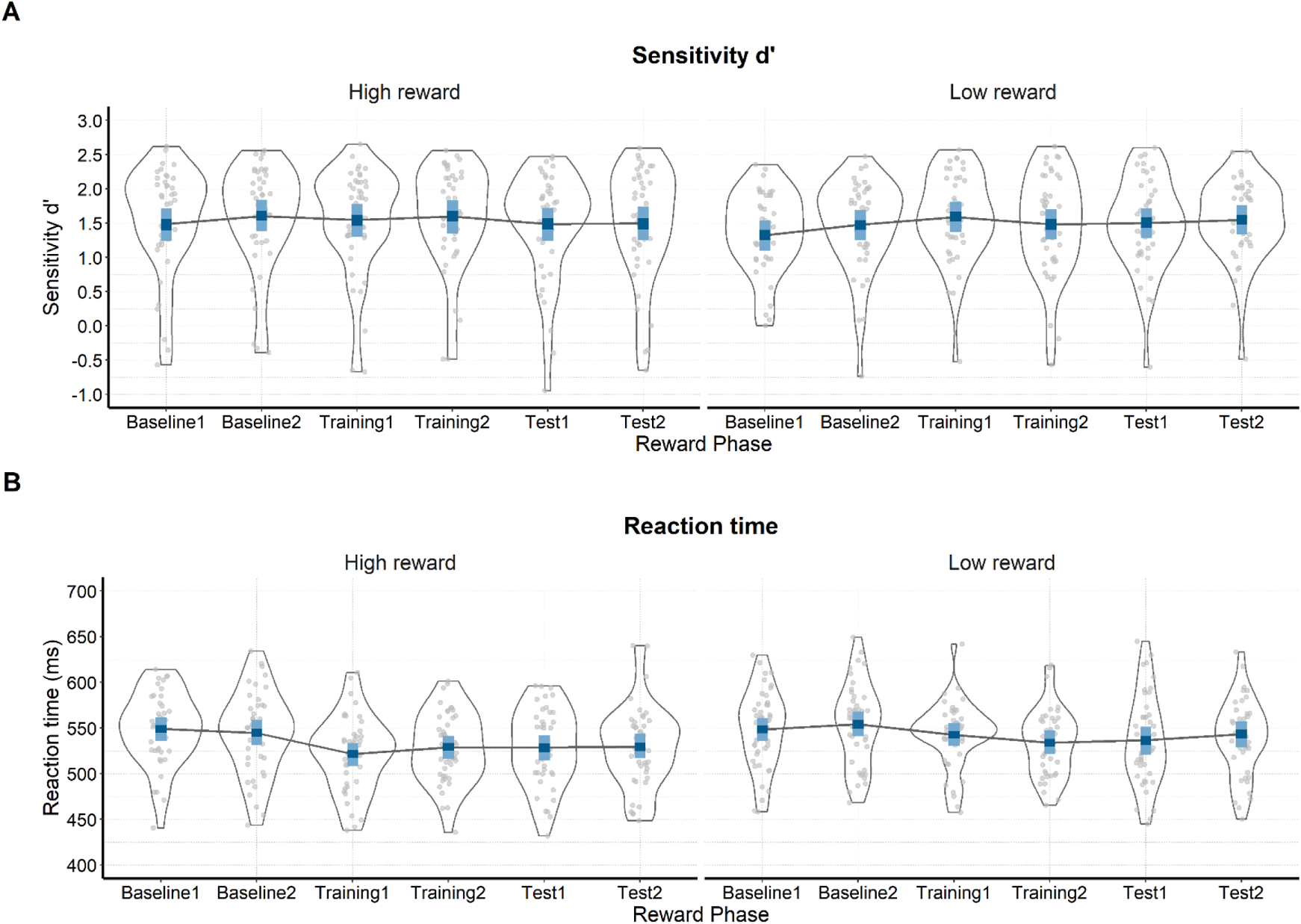
**Raw and modelled behavioral data in each half of each phase of the experiment.** On both plots raw participant data is represented with grey dots and their distribution. The winning model is presented in blue (dark blue – 50% HDIs and light blue – 95% HDIs). **A.** Sensitivity (d′) across the phases of the experiment for the conditions in which the attended color is linked to either high or low value. **B)** Reaction times (ms) in the six phases when the attended stimulus is related to high or low reward probability.

**Supplementary Table 4.** Means and 95% HDIs of sensitivity and reaction times across six phases of the experiment

| Reward phase | Value | Sensitivity (d') | Reaction times (milliseconds) |
| --- | --- | --- | --- |
| Baseline 1 | High | 1.48 [1.24, 1.71] | 548.86 [535.97, 561.35] |
| Baseline 1 | Low | 1.32 [1.09, 1.53] | 548.38 [535.83, 560.97] |
| Baseline 2 | High | 1.60 [1.38, 1.84] | 544.34 [531.22, 558.49] |
| Baseline 2 | Low | 1.47 [1.25, 1.69] | 553.98 [540.67, 567.69] |
| Training 1 | High | 1.54 [1.30, 1.78] | 521.42 [508.48, 533.66] |
| Training 1 | Low | 1.59 [1.37, 1.81] | 542.35 [530.05, 555.45] |
| Training 2 | High | 1.60 [1.35, 1.83] | 528.74 [515.92, 541.36] |
| Training 2 | Low | 1.48 [1.26, 1.70] | 533.91 [521.41, 547.24] |
| Test 1 | High | 1.49 [1.24, 1.72] | 528.64 [514.39, 542.24] |
| Test 1 | Low | 1.50 [1.28, 1.71] | 536.51 [520.49, 551.37] |
| Test 2 | High | 1.49 [1.25, 1.74] | 529.32 [516.53, 543.70] |
| Test 2 | Low | 1.55 [1.33, 1.76] | 543.01 [528.56, 557.28] |

## Acknowledgements

This work was supported by the Special Research Fund (BOF) of Ghent University [grant #01D02415 awarded to IG; grant # BOF14/PDO/123 awarded to AS], the Concerted Research Action Grant of Ghent University [grant number BOF16/GOA/017 awarded to EHWK], and the Biotechnology and Biological Sciences Research Council [BB/P002404/1 awarded to SKA]. The funding sources were not involved in the study design; collection, analysis, and interpretation of data; writing of the report; and decision to submit the article for publication.

We would like to thank Prof. Gilles Pourtois for his help with conceiving the study and for the very useful discussions of the results. Further, we thank Gilles for all of the materials he provided for this study. We would also like to thank Dr. Ladislas Nalborczyk for discussions about statistical analyses of the data, Prof. Ruth Krebs for her comments on a previous version of the manuscript, and Dr. Inez Greven for help with data collection.

## Author contributions

Author contributions are coded according to the CRediT taxonomy (Allen, Scott, Brand, Hlava, & Altman, 2014).

**Conceptualization**: IG, AS, EHWK, SKA. **Data curation**: IG, AS. **Formal analysis**: IG, AS. **Funding acquisition**: IG, AS, EHWK, SKA. **Investigation**: IG, AS. **Methodology**: IG, AS, SKA. **Project administration**: IG, AS. **Resources**: EHWK, SKA. **Software**: SKA, IG, AS. **Supervision**: AS, SKA. **Validation**: IG, AS. **Visualization**: IG, AS. **Writing – original draft**: IG, AS. **Writing – review & editing**: IG, AS, EHWK, SKA.

## Data availability

Raw and pre-processed data, materials, and analysis scripts are available at: https://osf.io/kjds3/.

## Software for data visualization and analysis

Visualization and statistical analyses were performed using R v3.4.4 (R Core Team, 2017) via RStudio v1.1.453 (RStudio Team, 2015). We used the following packages (and their respective dependencies):

- data manipulation: tidyverse v1.2.1 (Wickham, 2017);
- statistical analyses: Rmisc v1.5 (Hope, 2013), brms v2.3.1 (Bürkner, 2016);
- visualization: cowplot v0.9.2 (Wilke, 2016), yarrr v0.1.5 (Phillips, 2016), viridis v0.5.1 (Garnier, 2018), eegUtils v0.2.0 (Craddock, 2018), BEST (Kruschke & Meredith, 2017);
- report generation: pacman v0.4.6 (Rinker & Kurkiewicz, n.d.), knitr v1.20 (Xie, 2018);

## Footnotes

1 Throughout this manuscript we use the terms ‘attended’ and ‘unattended’ to refer to the explicit instructions which participants received prior to each trial.

2 Due to the simultaneous estimation of group-level and participant-level parameters, multilevel models display a property called shrinkage. In brief, estimates that strongly deviate from the mean (e.g., a participant performing the task much worse than the average of the total sample) will be pulled toward the group mean (McElreath, 2016). This advantageous property prevents extreme values from having large effects on the results.

